# Blood volume sensitive laminar fMRI with VASO in human hippocampus: Capabilities and biophysical challenges at clinical 7T scanners

**DOI:** 10.1101/2025.08.25.672075

**Authors:** Khazar Ahmadi, Stephanie Swegle, Sriranga Kashyap, Antoine Bouyeure, Peter Bandettini, Nikolai Axmacher, Laurentius (Renzo) Huber

## Abstract

Sub-millimeter resolution functional magnetic resonance imaging (fMRI) at ultra-high field (≥ 7T) has offered an unprecedented opportunity to probe mesoscopic computations at a columnar or laminar level. However, its application has been primarily restricted to the neocortex. Inferior brain regions, particularly the hippocampus (HC), are challenging targets for laminar fMRI. Recent developments in acquisition methods have shown the feasibility of laminar recordings in the HC using gradient-echo blood oxygenation level-dependent (BOLD) contrast. Nonetheless, the spatial specificity of the BOLD signal is compromised by the draining veins’ bias. Cerebral blood volume (CBV)-sensitive sequences including vascular space occupancy (VASO) have emerged as a promising augmentation tool to capture the vein-free laminar activity. Yet, its feasibility in the HC is unclear and challenged by methodological constraints. Here, we optimized VASO to mitigate the macrovasculature contribution in HC. By evaluating a series of advanced acquisition strategies tailored to HC, we obtained improved VASO signal quality with minimal artifacts. The optimized protocol was further validated with an autobiographical memory task. Our findings show that combining the high detection power of gradient-echo BOLD with the vein-free VASO contrast allows for differentiation between neural activity-related BOLD signals and those biased by draining veins. These results demonstrate the feasibility of submillimeter VASO acquired with conventional 7T scanners in the HC to map the circuit-level mechanisms of memory retrieval across HC subfields, laying a foundation to investigate the microcircuitry of HC-driven complex cognitive functions and their alterations in neurodegeneration.

## Introduction

Laminar fMRI is a rapidly growing field that enables the dissociation of feedforward and feedback projections, each targeting distinct cortical layers (1). This offers an opportunity to examine the mesoscopic organization of the brain at higher precision, facilitating the integration of microscopic insights from animal and postmortem studies with macroscopic findings from behavioral and conventional neuroimaging research in humans (1–4). Despite its increasing popularity, the vast majority of laminar fMRI studies have focused on the neocortex (5–11) while the underlying microcircuitry of allocortex, especially the hippocampus (HC) has been sparsely studied using this approach (12,13).

The HC, situated at the apex of the ventral visual stream, plays a key role in complex cognitive functions including memory and navigation. Although these functions have been well characterized at the macroscale, predominantly using conventional fMRI with spatial resolution of approximately 2 to 3 mm (14–16), their circuit-level substrates are poorly understood. This is partially due to signal drop-out associated with its proximity to air cavities that distort the magnetic field (12,13,17). Further, the increased distance between radiofrequency (RF) coil elements and lower brain areas including the HC combined with a large matrix size and shorter T2* values required for data acquisition lead to lower signal-to-noise ratio (SNR), making it difficult to capture the HC at ultra-high field (18). Additionally, the cytoarchitectonic laminar structure of the HC and its macrovasculature differ markedly from those of the neocortex (19), posing additional hurdles for the analysis of laminar fMRI data in this subcortical region. Unlike the neocortex with a canonical six-layered structure, the HC exhibits a three-layered laminar architecture arranged in a radial fashion following its curvature, similar to layers of an onion (13). Importantly, HC is not a homogenous structure, rather it comprises multiple subfields including subiculum, dentate gyrus (DG), and four cornu ammonis areas (CA1 to CA4), with CA4 also referred to as the hilus of the DG (20,21). These subfields are interconnected with neocortical areas via the entorhinal cortex (ERC). This is mediated by two major, anatomically distinct circuits (Figure.1A), the trisynaptic pathway which links superficial layers of ERC to DG and then to proximal apical dendrites of CA3 and subsequently to CA1 via the Schaffer collaterals; and the monosynaptic (temporoammonic) pathway that directly couples ERC to superficial layers of CA1 at distal apical dendrites in stratum radiatum lacunosum moleculare (SRLM). Within CA3, pyramidal neurons are interconnected through a recurrent autoassociative network toward stratum radiatum (22, 23). In contrast, the HC output originates from deep layers of CA1 at stratum pyramidale and stratum oriens and is later funneled to deep layers of ERC (12,17,24,25).

**Figure 1.**
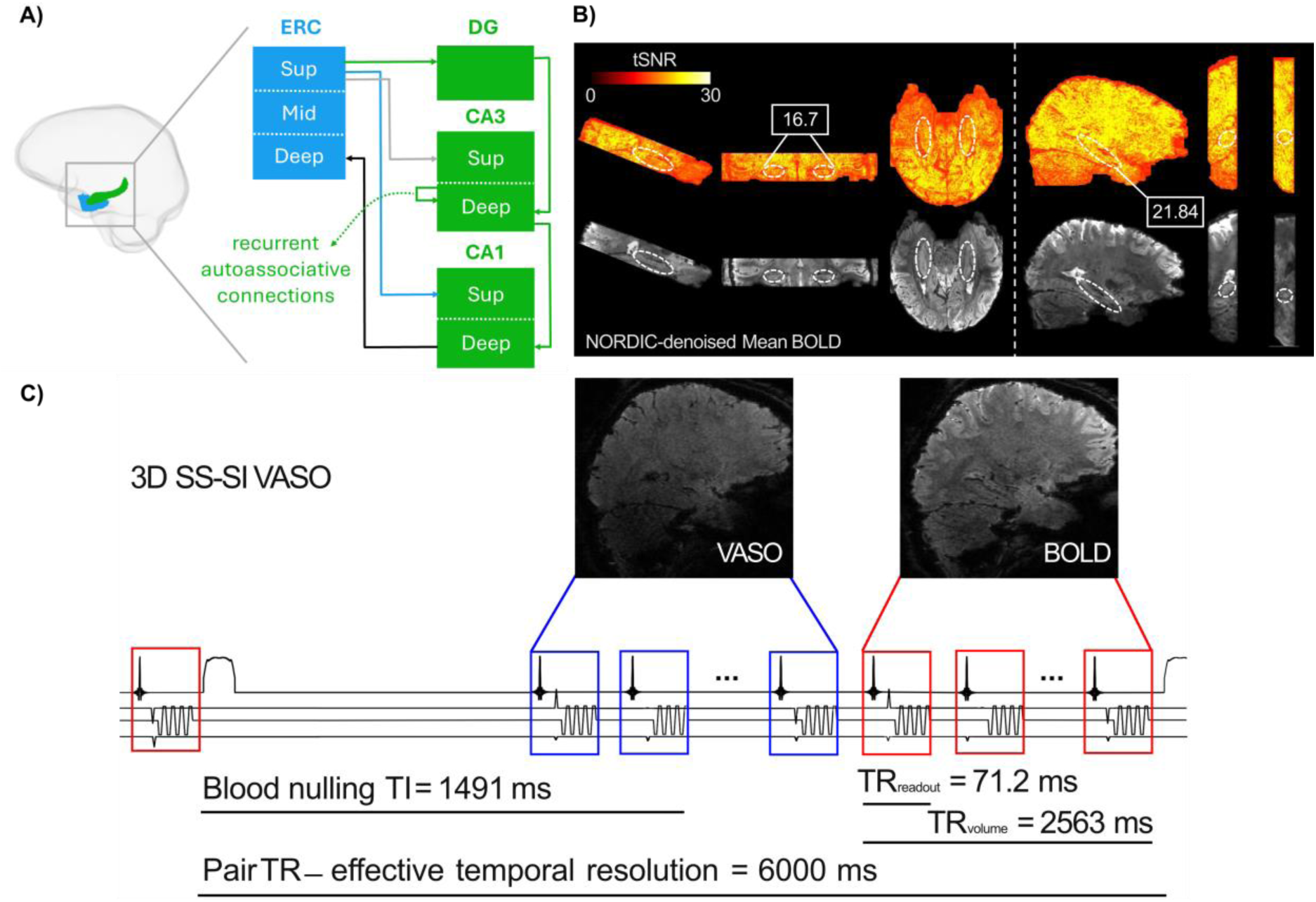
Overview of hippocampal projections and VASO optimization procedure. **A)** Schematic illustration of HC-ERC microcircuitry. The trisynaptic pathway (green) includes projections between ERC, DG, CA3, and CA1 while the monosynaptic (temporoammonic) pathway (blue) comprises direct connections from ERC to CA1. There are also direct connections between ERC and CA3, depicted in grey. The HC output (black) targets deep layers of ERC. The dashed green arrow represents autoassociative connections between CA3 neurons. ERC = Entorhinal Cortex, HC = Hippocampus, Sup and Mid = superficial and middle layers, respectively. The SRLM (not shown here) lies within the superficial compartment in CA1. **B)** tSNR maps and mean BOLD images from one participant, acquired using the reference 3D BOLD-based EPI sequence (left panel) with bilateral HC coverage and imaging slab oriented parallel to the HC long axis. The right panel shows the resulting images after adjustment of multiple parameters including FOV reduction, targeting left HC in sagittal orientation in the same participant. The optimization led to improved tSNR, as shown in the white box. This improvement is expected to generalize across participants since the adjustments systematically reduce artifacts caused by scanner hardware induced EPI phase inconsistencies. Dashed circular markers indicate approximate location of HC. **C)** Schematic depiction of HC-tailored 3D SS-SI VASO sequence protocol that was developed by further refining the acquisition parameters in B right panel. Note that ∼ 500 ms was dedicated to slice-wise fat suppression modules, increasing the ultimate volume TR to 3000 ms and effective temporal resolution to 6000 ms. TR = repetition time, TI = inversion time, SS-SI = slice-selective slab-inversion.

Despite these challenges, recent advancements in gradient-echo sequences with blood oxygenation level-dependent (GE-BOLD) contrast and echo planar imaging (EPI) readouts (26) alongside the availability of new analysis approaches and software packages have driven an increasing interest in mapping the laminar organization of HC (13,27). While the BOLD contrast provides a relatively high SNR (28,29), it is prone to venous drainage effects. This is particularly pertinent in HC where the blood can drain from either the inner surface (i.e., superficial layers) or outer surface (i.e., deep layers) (19). The contribution of large vessels to laminar BOLD responses has been extensively studied in the neocortex and several physiological models have been developed to account for venous bias (30–32). However, such models are currently lacking for the HC whose vasculature is substantially different from the neocortex. This warrants the application of alternative acquisition approaches such as blood volume-weighted laminar fMRI with vascular space occupancy (VASO) that hold a great potential to augment the high sensitivity of BOLD with an additional measure of vein-free laminar-specific activation (33). VASO has been successfully implemented to resolve laminar activity across various cortical areas (5,34–36). Nonetheless, its application in HC remains unexplored. Building on a previous laminar fMRI study with BOLD contrast (13), we sought to develop an effective VASO protocol to map the laminar organization of HC across typical 7T scanners. Here, we first outline a series of advanced acquisition strategies to mitigate the physiological and methodological challenges of VASO in the HC that are essential for optimizing an HC-tailored imaging protocol. We then present the results of a validation study to assess the laminar profiles of the HC subfields during an autobiographical memory paradigm (13,37,38). Our findings establish the feasibility of mesoscale VASO recordings in HC, providing groundwork for future studies to decipher the circuit-level mechanisms of high-level cognitive processes in HC.

## Methods

### Participants

Thirteen healthy volunteers (6 females, mean age = 30.7 years) were recruited from the National Institutes of Health (NIH) community after providing written informed consent according to procedures approved by the NIH Institutional Review Board. A total of 18 scanning sessions were conducted where ten participants each completed a single session, while three participants took part in multiple sessions (two participants completed two sessions each, and one participant completed four sessions). The study was conducted in two parts. The first part focused on optimizing the VASO sequence, during which 3 participants underwent 4 sessions. The second part, aimed at validating the optimized sequence with an autobiographical memory paradigm, involved 11 participants across 14 sessions. Data from five individuals were excluded from the 2^nd^ part due to experimental errors (stimulus onset not synchronized with the sequence trigger, N = 2) or excessive motion artifacts (N = 3), leaving six usable datasets acquired over 9 sessions. Although no statistical tests were used to pre-determine the required sample size, the number of recruited individuals aligns with a previous BOLD-based laminar fMRI study utilizing the same task (13).

### Part 1: Developing an HC-tailored VASO protocol

Three 7T SIEMENS scanners, two identical MAGNETOM TERRA and a classic 7T Plus systems (SIEMENS Healthineers, Erlangen, Germany) were used depending on the availability of free timeslots. Each scanner was equipped with a 32 receive channel head coil (NOVA Medical, Wilmington, MA). All scanners operated with the same sequence version of the same software baseline VE12U, utilizing the same gradient strengths. Third-order shims were unplugged to reduce gradient resonance artifacts known as Fuzzy Ripples that are associated with magneto-mechanical coupling between third-order shims and read-out gradients (18). Additionally, the parallel transmission (pTX) system was used because of its larger transmit coverage in lower brain areas (18). To quantify temporal signal-to-noise ratio (tSNR) in the HC, we first employed a reference 3D GE-EPI sequence with BOLD contrast (26) in line with previous studies (13,27). Briefly, the key parameters were TR|TE = 2500|28.4 ms, voxel size = 0.8 x 0.8 x 0.8 mm^3^, FOV = 192 mm, GRAPPA = 4, base resolution = 240, FA = 14°, bandwidth = 1096 Hz/px, partial Fourier = 7/8, phase-encoding direction = AP and 40 slices where imaging slab covered bilateral hippocampi and was placed parallel to their long axis. To boost the tSNR, the imaging slab was positioned in sagittal orientation with unilateral HC coverage (see Figure.1B) and several acquisition parameters of the 3D GE-EPI sequence (26) were adjusted as following: FOV = 180 mm, GRAPPA = 3, base resolution = 226, partial Fourier = 6/8, bandwidth = 1106 Hz/px, TR|TE = 2664|21.8 ms, FA = 18°. Leveraging this optimized acquisition setup, a 3D slice-selective slab-inversion (SS-SI) VASO sequence was implemented to acquire concomitant VASO and BOLD images in an interleaved fashion (33). VASO uses an inversion recovery pulse to null the blood signal enabling the measurement of cerebral blood volume changes through the residual tissue signal (39,40). A major methodological challenge of VASO in the HC is the relatively short arterial arrival time (41) which can lead to inflow of fresh (non-inverted) blood during VASO read-out, manifesting as bright spots in the acquired data. To implement a VASO protocol tailored to HC, we adjusted a wide range of parameters including read-out and volume TR, TE, FA, TI1, TI2, inversion delay, RF power scale and RF Bandwidth Time product (BWTP). The optimum set of parameters were determined with respect to tSNR and minimal inflow contamination in HC (see Part 2 and Results). Following the optimization of the VASO sequence, one participant was re-scanned with a single transmit (1sTX/32Rx) head coil allowing for direct comparison with pTX coil. For anatomical reference, whole-brain T1-weighted images were acquired using 3D Magnetization Prepared 2 Rapid Acquisition Gradient Echo (MP2RAGE) sequence (42) at an isotropic resolution of 0.75 mm^3^ with TR|TE = 4550|1.94 ms, TI1|TI2 = 840|2370 ms, FA1|FA2 = 5°|6° and matrix size = 240 x 320 x 320.

### Part 2: Sequence validation with an autobiographical memory paradigm Experimental task

The autobiographical memory paradigm was implemented in a block design using PsychoPy (43), adapted from previous studies (13,37,38). It consisted of two conditions: i) autobiographical memory retrieval and ii) math with 15 trials per condition. The math trials served as a baseline since mental arithmetic primarily activates the parietal cortex (44–46) and does not rely on episodic memory and HC function (38). Each trial lasted for 18 seconds followed by an inter-trial interval of 12 seconds during which a central blue square was displayed on a white background. During memory trials, participants were presented with general word cues e.g., “beach”. They were instructed to recall a cue-related event from their personal past and re-experience it as vividly as possible for the remainder of the trial. To ensure the HC involvement, participants were required to retrieve recent memories (no older than 2 to 3 years) as remote episodic memories may be more strongly represented in neocortical regions (47,48). In the math condition, participants solved a simple addition or subtraction problem, for example, “74 – 35”. Upon finding a solution, they were instructed to iteratively add 3 to their answer e.g., (39 + 3 + 3 + 3 …) until the end of the trial. Prior to the scanning, participants were thoroughly briefed on the experimental procedure. Each participant completed a minimum of 3 runs, with each run consisting of 15 trials featuring a unique set of word cues and math operations. The stimuli were presented on a rear-projection screen located at the magnet bore and viewed through an angled mirror.

### Data acquisition

Using the optimized protocol, we acquired concurrent VASO and BOLD data from 6 participants across 9 sessions with the same scanners described in part 1. Each session consisted of three runs. Toward the end of each session, participants were asked via intercom whether they were willing to complete an optional fourth run. As a result, four runs were obtained in three participants. A few runs in some participants were discarded due to excessive motion artifacts, thresholded as maximum displacement of 2mm due to rotation or translation. Detailed information on the number of acquired sessions and analyzed datasets per participant is provided in Supplementary Table1. The acquisition parameters were as follows: Inversion-recovery 3D-EPI (26,49) with isotropic voxel size of 0.8 mm^3^, 36 slices, FOV = 180 mm, read-out TR | volume TR | TE = 71.2, 6000, 23.90 ms, PE-direction = AP, variable flip angles with reference (last) nominal value of FA = 35°, base resolution = 226, bandwidth = 1106 Hz/pixel, GRAPPA = 3, partial Fourier = 6/8, RF BWTP = 12, RF power scale = 3, , TI1 = 1491 ms. Each functional run comprised 308 time points, corresponding to 154 pairs of interleaved VASO and BOLD volumes with a total duration of 15.4 minutes. In addition, anatomical data were acquired in all participants with identical parameters specified in part 1. When possible (N = 5 out of 6), an auxiliary fMRI scan with opposite phase-encoding direction i.e., PA, was acquired to facilitate correction of susceptibility-induced distortion artifacts.

## Data analysis

### Anatomical data processing

The unified image (UNI) of the MP2RAGE sequence was first preprocessed using Presurfer (50) to remove background noise by utilizing a bias-corrected image corresponding to second inversion time (INV2). The resulting denoised image was then inputted to HippUnfold package (51), executed via Singularity, to automatically segment HC subfields and delineate its surface boundaries (Supplementary Figure.1). Although T2-weighted structural images are generally favored for HC segmentation owing to superior contrast between HC grey matter (GM) and the SRLM tissue, recent advances in HippUnfold enable robust and accurate HC segmentation on T1-weighted data (51,52). The denoised T1 image was further skull-stripped using FSL brain extraction tool (BET; 53) and subsequently segmented with FSL FAST (54) to classify three tissue types corresponding to GM, white matter (WM) and cerebrospinal fluid (CSF).

### Functional data preprocessing

All functional runs were sorted by contrast yielding separate VASO and BOLD time series, each comprising 154 volumes (55). The first two volumes of each contrast were discarded in every run to allow magnetization to reach a steady state. Of the 36 acquired slices, three were removed from the outermost edge of the acquisition slab due to distortion artifacts. Noise Reduction with Distribution Corrected Principle Component Analysis (NORDIC-PCA; 56–58) was applied for each contrast separately to boost the inherently low SNR of the laminar fMRI data through thermal noise suppression. The last two volumes, that are typically used for estimation of the noise threshold in NORDIC, were also excluded resulting in a total of 150 volumes per contrast. Motion and distortion correction were performed using Advanced Normalization Tools (ANTs). Following a rigid-body motion correction, a temporal mean image was generated for time series of VASO and BOLD contrasts and their corresponding reversed phase-encoding data separately. Afterwards, the mean image of the reversed phase-encoding data was rigidly aligned to the motion-corrected mean image of each contrast’s time series. The two images were subsequently fed to antsMultivariateTemplateConstruction2 (59) to perform a ‘meet-in-the-middle’ registration using ANTs symmetric normalization (SyN) algorithm to obtain distortion-corrected mean images for each contrast. Next, all transformation and distortion-corrected warps were concatenated and applied using antsApplyTransforms with Lanczos interpolation. Note that distortion correction was applied in five out of the six datasets as opposite phase-encoding data was not available in one participant (see Data acquisition). The preprocessed VASO data were further corrected for BOLD contamination using the LN_BOCO program in LayNii (60). Following preprocessing, the BOLD-based fMRI data were aligned to the T1 image using the AFNI script ‘align_epi_anat.py’. The mean VASO image was first meticulously aligned to the T1-weighted image using ITK-SNAP (61), utilizing its capability of allowing careful manual interventions to ensure high-quality alignment in the presence of EPI artifacts in some parts of the FOV that can confuse fully automatic registration approaches. The resulting transformation matrix was then applied to all VASO time series with fifth order B-spline interpolation. Further, to quantify physiological noise in the T1-aligned VASO and BOLD time series, anatomical Component Correction (aCompCor; 62,63) was applied. Specifically, five principal components were extracted separately from WM and CSF masks, resulting in 10 regressors in total. Prior to aCompCor implementation, the whole-brain WM and CSF masks were cropped to match the co-registered functional data and eroded using the FSL tool ‘fslmaths’ with a Gaussian kernel of σ = 0.8 mm to ensure that no HC voxels were included in the anatomical masks. All stages of preprocessing were subject to careful visual inspection for quality control. Moreover, tSNR maps were computed in all functional time series to quantitatively assess the data quality.

### Extraction of laminar profiles in HC subfields

In line with previous studies (13,27), the co-registered VASO and BOLD time series were sampled as a function of depth using a custom MATLAB code. In short, GM signal in each HC subfield was sampled into 20 equidistant bins spanning the vertices of the HC inner and outer surfaces, that were generated by HippUnfold (51). To account for the pronounced curvature of the HC, the mid-thickness depth, also computed via HippUnfold representing the center of GM in the HC, was designated as a reference landmark for the sampling algorithm. In subiculum, CA1, and CA2, the sampling was further extended by an additional 10 bins beyond the inner surface to encompass the SRLM. The sampling was not applied to DG/CA4 due to the absence of a well-defined inner/outer surface boundary in these areas (see Supplementary Figure.1).

### General linear model (GLM) analysis

GLM analysis was performed using Statistical Parametric Mapping (SPM12; Wellcome Trust Centre for Neuroimaging, London, UK) to separately estimate the VASO and BOLD activation maps for each stimulus condition. No spatial smoothing was applied. The fMRI data underwent high-pass filtering with a default cut-off of 128s and were convolved with a canonical hemodynamic response function. A design matrix was created, incorporating two regressors of interest i.e., autobiographical memory and math conditions, as well as 16 nuisance regressors including 6 motion estimates and 10 principal components derived from aCompCor. For each participant, a primary contrast of interest was calculated as the difference between beta-estimates of memory and math conditions. Statistical maps were computed across the entire slab volume of the fMRI data. To mitigate the problem of multiple comparisons with the increased number of voxels at high spatial resolutions, and enhance anatomical specificity, a GM mask cropped to the functional slab coverage was applied during cluster-level inference.

For group-level analysis, a study-specific anatomical template with an isotropic spatial resolution of 0.75 mm^3^ was generated from the denoised T1-weighted images of all participants using the ANTs multivariate template construction algorithm. The statistical maps from first-level GLM analysis were subsequently co-registered to this template using the transformation matrices obtained during template construction.

The GLM analysis was also conducted on depth-dependent VASO and BOLD responses across HC subfields. Following the approach outlined by (13), the laminar time series were first z-transformed using the mean and standard deviations computed from the volumes corresponding to math condition.

## Results

### Part 1: Developing an HC-tailored VASO protocol

A schematic illustration of the optimized protocol is presented in Figure.1C and a detailed list of the protocol parameters is available in the following GitHub repository: https://github.com/layerfMRI/Sequence_Github/tree/master/Hippocampus.

### Mitigating methodological and physiological challenges of VASO in HC

EPI quality was improved through i) read-out along the z-axis to minimize short-term eddy currents, ii) bandwidth adjustments, and iii) disconnecting third-order shim to decrease inductive coupling (Figure.2A; see also ref.18). To evaluate the inversion efficiency of inflowing blood in the HC, a series of adiabatic RF pulses with varying power scales were applied. The aim was to determine whether inflow effects remain confined to upstream vascular compartments or propagate downstream into the HC microvasculature. Two scenarios were considered: i) if no change in inflow-related signal was observed across RF power levels, this would imply that inversion inefficiency affects smaller vessels downstream, including the HC microvasculature. ii) If inflow effects decrease with stronger RF power, this would suggest that inflow effects are largely restricted to large upstream vessels, with minimal impact on the HC microvasculature. The results in Figure.2B support the 2^nd^ scenario. Inflow-related signal increased at lower RF powers and was attenuated with stronger inversion (optimized RF power scale = 3), indicating that inflow effects are mostly limited to larger vessels. Residual inflow signal in these vessels is not a concern, as they do not exhibit task-related functional changes and maintain constant flow between rest and task conditions. Upon optimizing inversion pulse parameters, no major inflow-compromised VASO signal was detected in HC. Notably, inflow modulations were comparable between the HC and some posterior neocortical regions (Figure.2B).

**Figure 2.**
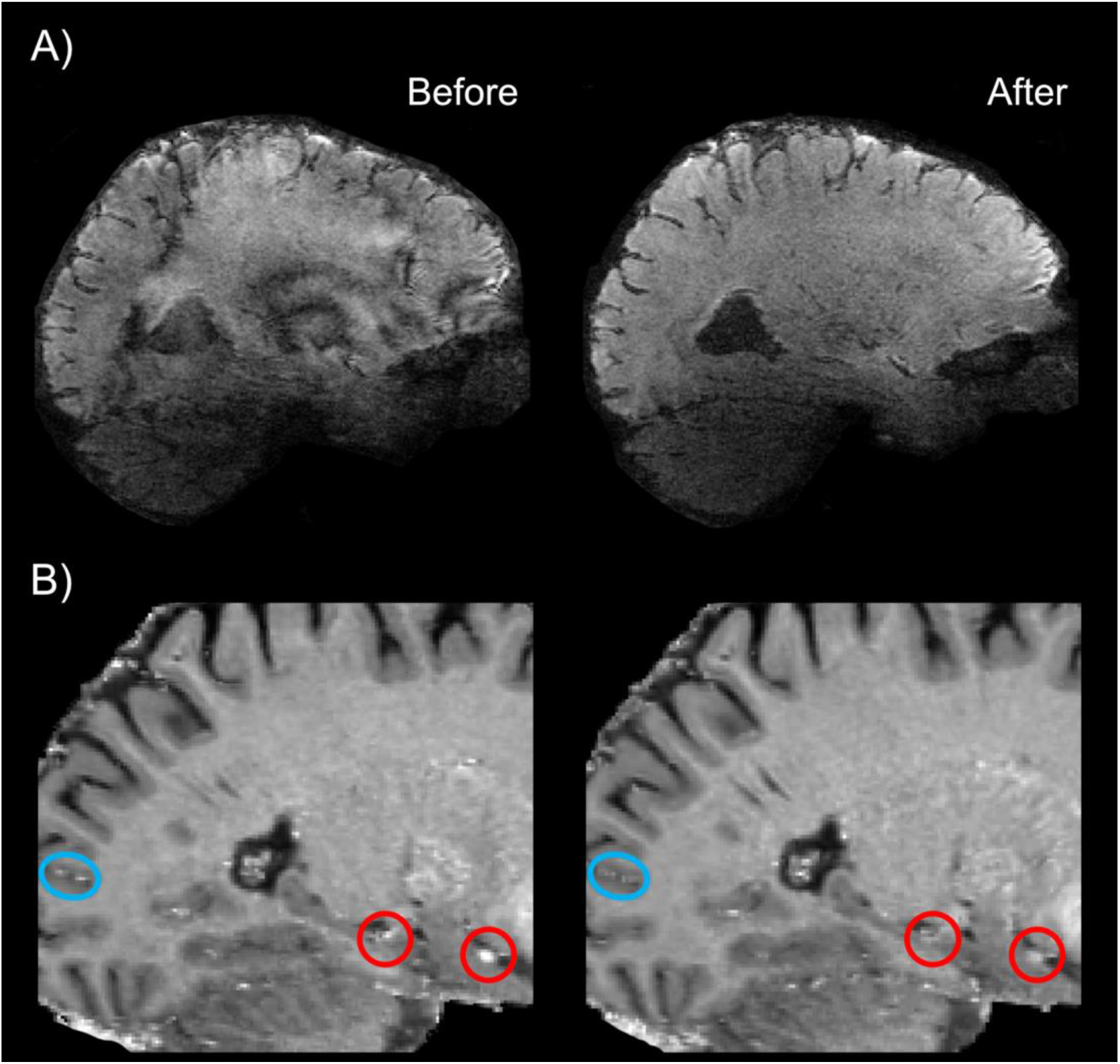
Characterization and mitigation of gradient-related and inflow artifacts in VASO-based laminar fMRI in HC. **A)** Bandwidth adjustments, disconnecting third-order shim and read-out along the z-axis collectively mitigated the EPI phase errors. **B)** Inflow effects were assessed using a series of adiabatic RF pulses of varying power. Blue and red circles indicate inflow-related signal modulations in a posterior neocortical region and the HC as well as a nearby MTL region, respectively. The bottom right panel demonstrates that increasing the RF power of the inversion pulse decreases inflow-related signal modulations in large vessels outside of GM. This suggests that inflow effects are not expected in downstream microvessels of the GM tissue. EPI = echo planar imaging, HC = hippocampus, MTL = medial temporal lobe, GM = grey matter.

### Sequence stability across scanners and comparison of coil performance

Three conventional 7T scanners (see Methods) were used in this study, allowing us to investigate the robustness of the HC-tailored VASO protocol across scanners. The mean tSNR in the HC as well as the mean image quality for both VASO and BOLD contrasts were comparable across different scanners, indicating that the performance of the optimized sequence is independent of scanner-specific factors (Figure.3A). Due to the unequal number of data points per scanner, statistical testing was not conducted to assess between-group differences to avoid potential bias. Additionally, in one participant who was scanned using both sTX and pTX coils, HC-tSNR values were compared between coil types. As shown in Figure.3B, the sTX coil yielded slightly higher tSNR for both BOLD and VASO contrasts in this participant, suggesting a potential advantage for HC-focused laminar applications.

**Figure 3.**
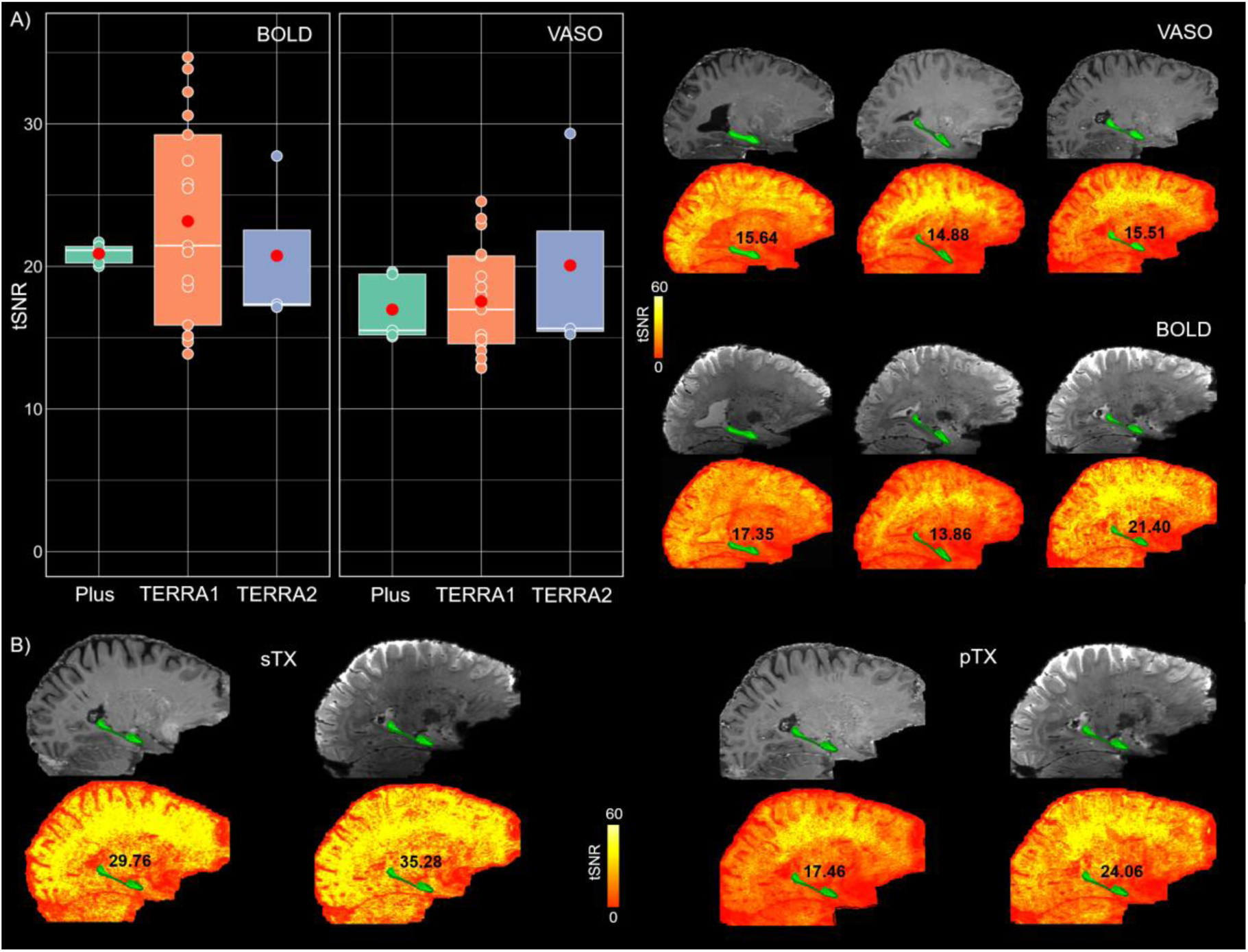
Consistency of tSNR in HC across scanners and head coils. **A)** Mean HC-tSNR values for BOLD and VASO contrasts across two identical MAGNETOM TERRA and one classic 7T Plus scanners (N_TERRA1_ = 17, N_TERRA2_ = 3, N_Plus_ = 5). In the box plots, the white solid line denotes the median, the red circle represents the group mean and the remaining scatter points reflect the individual data per scanner. The top right panel displays mean VASO and BOLD images along their corresponding tSNR maps in three randomly selected participants whose data are also shown in the box plots. Average HC-tSNR values are shown on the maps next to the anatomical location of HC, highlighted with a green mask. **B)** Comparison of mean BOLD and VASO images together with their respective tSNR maps between sTX and pTX coils in the same participant, indicating higher values for the sTX coil. HC = hippocampus, tSNR = temporal signal-to-noise ratio.

### Part 2: Sequence validation with an autobiographical memory paradigm Disentangling brain activations between memory and math trials

First-level GLM analysis with a cluster extent threshold of 20 voxels revealed predominant activation in parietal regions during math operations whereas memory retrieval elicited greater activity in the frontal cortex, parahippocampal cortex and the HC in all participants. Importantly, the overall activity patterns were consistent between VASO and BOLD contrasts (see Figure.4A). To investigate subfield-specific effects, signal changes were further examined within significant subject-specific activation clusters in the HC (Figure.4A). Specifically, for each participant, the mean signal change was extracted from portions of the cluster that overlapped with their anatomically defined HC subfields. As shown in Figure.4B, the mean signal change for both imaging contrasts was comparable in all subfields with no statistical difference detected by the Friedman test (VASO: χ²(5) = 2.63, p = 0.75; BOLD: χ²(5) = 4.76, p = 0.44), suggesting a uniform engagement of HC subfields during autobiographical memory retrieval. Given the small sample size, the non-parametric Friedman test was used instead of parametric alternatives such as repeated-measures ANOVA.

**Figure 4.**
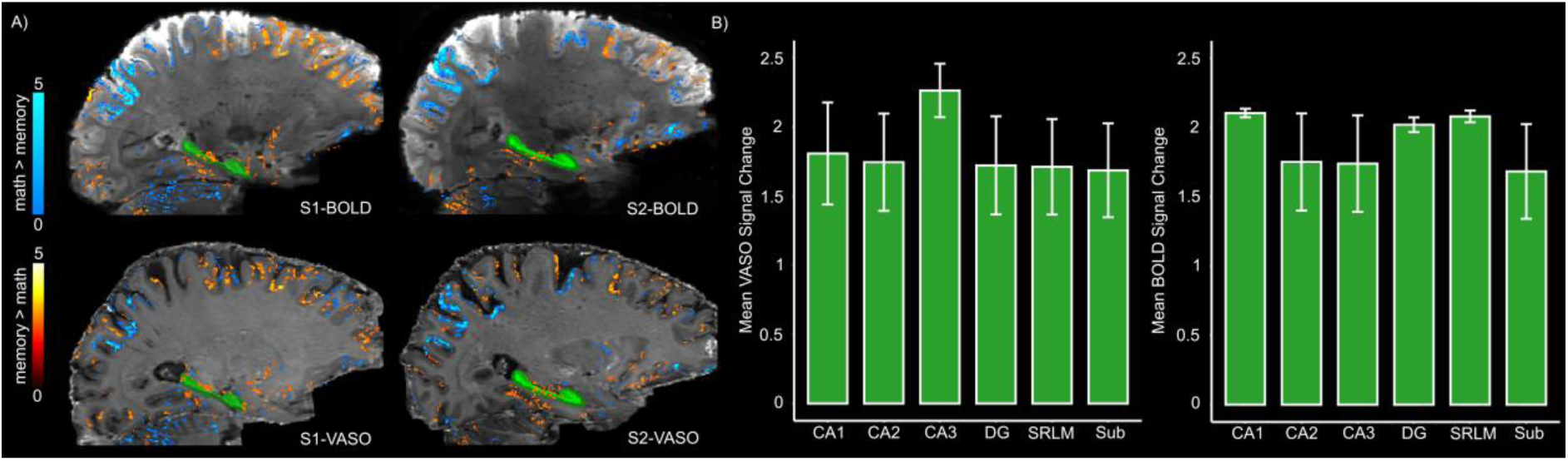
Brain activation maps and average signal changes across HC subfields for BOLD and VASO contrasts. **A)** Representative activation patterns from two participants (S1 and S2) during memory and math trials overlaid on the mean BOLD (top) and VASO (bottom) images (p < 0.05, uncorrected). The green mask indicates HC. **B)** Mean signal change for VASO and BOLD (left and right panels, respectively) across subfields derived from significant clusters within the HC for memory > math (as shown in panel A). The error bars indicate standard error of the mean (SEM). Note that here ‘DG’ represents a merged ROI encompassing both the dentate gyrus and CA4.

Similar activation maps (nifti data available on Zenodo) were observed at the group level although the effects were less pronounced in the VASO dataset. The similarity of the observed activation patterns during memory retrieval and math operations to previously reported activation profiles (37,38,44,64) further supports the validity of the optimized sequence.

### Laminar profiles of HC subfields for memory vs math trials

Figure.5 illustrates the average depth-dependent profiles of HC subfields across all participants for the z-transformed memory > math contrast (see Methods and ref.13) in BOLD and VASO data. In subiculum and CA1, the laminar BOLD signal exhibited a peak at the SRLM adjacent to the inner surface, corresponding to superficial layers, followed by a decreasing pattern towards the outer surface. In CA2, the BOLD signal showed a modest elevation at SRLM, a dip in the inner surface and a subsequent increase at the outer surface. In contrast, CA3 displayed a largely monotonic increase in the BOLD signal across depths. Crucially, the increase in BOLD-driven laminar profiles across HC subfields aligns with the expected draining venous bias based on neuroanatomical evidence and a recent BOLD-based laminar fMRI study (19,13), i.e., a more pronounced effect towards the SRLM/inner surface in the subiculum and CA1 and at the outer surface in CA2 (see Figure.5). Interestingly, the VASO signal demonstrated a primarily inverse laminar trend relative to BOLD, manifesting a decreasing pattern at the SRLM/inner surface of subiculum and CA1 as well as the outer surface of CA2. In CA3, however, the laminar profiles of BOLD and VASO were comparable. Similar laminar profiles were also observed for non-transformed memory > math contrast (Supplementary Figure.2).

**Figure 5.**
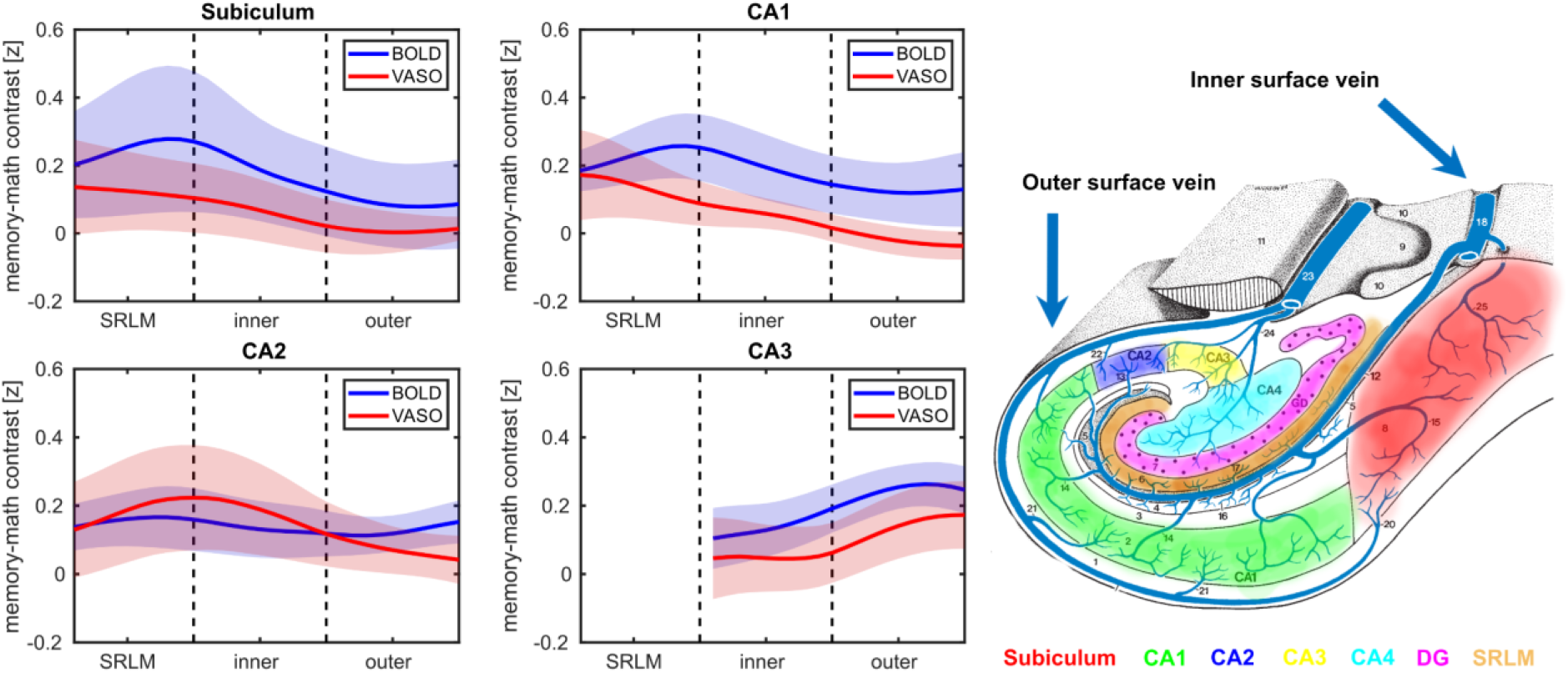
Laminar profiles of subfields for memory vs math contrast along with HC vascular architecture. In subiculum and CA1, the BOLD signal profiles (blue) peak at the SRLM near the inner surface whereas in CA2 the BOLD increase is confined to the outer surface, likely reflecting subfield-specific draining veins’ bias. Conversely, the VASO signal (red) shows the opposite pattern in those subfields at the same depth. Shaded areas indicate SEM. Note that the laminar profiles are not estimated in DG and CA4 due to lack of a clear boundary between the inner and outer surfaces. The right panel displays a schematic illustration of HC venous supply, adapted from ref.19.

## Discussion

Despite significant progress in harnessing laminar fMRI to uncover the directionality of information flow in the neocortical areas (6–9,65,66), analogous investigations in non-neocortical regions and particularly the HC remain limited (12,13,27). To date, the available studies have examined depth-dependent activity across HC subfields using GE-BOLD contrast whose signal is weighted towards large venous vessels. To mitigate the macrovasculature bias, we aimed to develop an HC-tailored VASO protocol. Our findings provide evidence for successful implementation of mesoscale VASO in human HC by i) yielding sufficient signal quality with minimal artifacts across conventional 7T scanners and ii) enabling reliable measurement of laminar-specific signal modulations across subfields during an autobiographical memory task.

### VASO optimization in HC

The major innovation of this study lies in the acquisition strategies designed to address the methodological and physiological challenges of applying submillimeter VASO in the HC on conventional 7T scanners. The anatomical location of the HC renders it susceptible to B_0_ and B_1_ inhomogeneities, leading to more pronounced EPI artifacts and a subsequent decrease in SNR (12,13,17). Additionally, similar to other inferior brain areas, the HC is located far from RF coil elements and often requires a relatively large matrix size and short T2* values for acquisition (18), factors that further exacerbate the SNR loss. We used pTX system and optimized the EPI protocol to boost the tSNR by decreasing FOV, GRAPPA, base resolution, and partial Fourier while increasing the bandwidth, and made further adjustments to other sequence parameters (see Methods, and Figure1.B&C).

A prominent source of artifact in laminar fMRI is the appearance of Fuzzy ripples represented by low spatial frequency signal shadings (18). These artifacts are particularly severe in deep brain areas including the HC and are largely associated with eddy currents in ramp-sampling EPI. In SIEMENS whole-body scanners, these eddy currents are predominantly induced by inductive coupling with the third-order shim (67). To mitigate Fuzzy ripples and improve the EPI equality, we unplugged third-order shim prior to data acquisition, reoriented the EPI read-out along the z-axis and adjusted the bandwidth (Figure.2A). Inflow-compromised VASO signals were largely mitigated using adiabatic RF pulses optimized for inversion efficiency (see Figures. 1C and 2B). After adjusting RF power scale, inflow effects in HC were similarly modulated to those observed in posterior neocortical regions, suggesting that the effects are primarily driven by larger arterial vessels with minimal impact on the HC microvasculature. The optimized protocol showed strong reproducibility across three conventional SIEMENS 7T scanners (two identical TERRA and a classic 7T Plus), representing the most widely distributed 7T platforms globally, thereby underscoring its generalizability and practical utility. Future work might include further cross-scanner comparisons across a more generalizable range of scanner hardware and task implementations (68). The comparison of head coils revealed a modest SNR advantage in HC for both BOLD and VASO contrasts with the sTX coil (Figure.3B), which is consistent with reports on increased noise coupling of the lower row of receivers in the pTx coil (69). While the number of available datasets obtained with both coil types (N = 1) precludes any conclusion, the observed difference may reflect g-factor amplification in the pTX coil (70). Future studies are, thus, warranted to explore this effect systematically in a larger cohort.

### Functional validation using a memory task

To assess the sensitivity of the optimized sequence, we employed an autobiographical memory paradigm. In line with previous studies (13,37,38), we observed HC, parahippocampal and frontal activity during memory retrieval for both BOLD and VASO data. Notably, both imaging contrasts yielded consistent activation patterns (Figure.4A). Further investigation of BOLD and VASO signal modulation in significant clusters within the HC showed no differential activation across subfields, suggesting a coordinated engagement of HC during autobiographical memory retrieval. This is partially in agreement with the findings of a recent study (38) reporting no significant difference in the activation of CA1 to CA3, DG/CA4 and the subiculum. However, that study employed manual segmentation of the subfields and included pre/parasubiculum which showed higher signal change than all other subfields. In contrast, we used HippUnfold (51) to automatically segment the subfields that does not delineate the pre/parasubiculum area. This may account for the absence of a distinct subfield effect in our results. Collectively, the convergence of the observed activation maps between BOLD and VASO coupled with their correspondence to regions identified in prior studies (37,38,44,64) support the robustness of the optimized sequence in detecting task-related activity in the HC.

Laminar analyses revealed distinct activation profiles in each subfield. Specifically, the laminar BOLD responses in subiculum and CA1 showed a peak at the transition zone between SRLM and the inner surface while in CA2 a modest increase in BOLD signal was observed towards the outer surface. These findings are largely consistent with a recent BOLD-based laminar fMRI study (13) and the neuroanatomical evidence on hippocampal venous supply (19) suggesting a potential contribution of draining vessels to the observed BOLD signal. In contrast, VASO signal profiles exhibited an inverse trend at the SRLM/inner surface border of subiculum and CA1 as well as the outer surface of CA2, likely reflecting its sensitivity to microvascular CBV changes. In CA3, the laminar profiles of BOLD and VASO were comparable, both showing a peak at the outer surface (corresponding to deep layers). This could potentially demonstrate the contribution of trisynaptic pathway to autobiographical memory retrieval via pattern completion, a key computational function governed by CA3. This process involves the reactivation of an entire memory or episode from partial cues and is thought to rely on recurrent autoassociative connections in deep layers of CA3, likely at stratum radiatum and proximal apical dendrites (22,23,71), towards its outer surface. Altogether, these findings indicate that the optimized VASO protocol can resolve depth-dependent activity in the HC.

The present study has a few limitations that should be acknowledged. First, the sample size was relatively small (N = 13). This is particularly relevant for part 2, the validation study, where the GLM results and the laminar profiles were derived from six datasets reducing the statistical power of the analyses. The observed effect sizes in Figures 4 & 5 were relatively small, which may be partially attributed to the limited amount of usable data. Although we acquired at least 3 functional runs per participant, some runs were discarded due to motion artifacts (see Supplementary Table.1). Moreover, each run contained 150 volumes after preprocessing whereas in the previous BOLD-based laminar fMRI study (ref.13) 489 volumes were acquired per run, resulting in a larger magnitude of the depth-dependent signal. The concomitant acquisition of BOLD and VASO data in our protocol leads to a lower temporal resolution, limiting the number of volumes that could be acquired within a reasonable scan duration as extending the acquisition time to collect more volumes would likely compromise data quality due to motion. Further, we applied NORDIC denoising (56,58) to boost the tSNR of the data. However, recent reports suggest that NORDIC may attenuate task-relevant signal variance (72), potentially contributing to the modest effect sizes observed in our study. Lastly, we used the default canonical HRF from SPM12 whereas prior work has shown that subcortical regions may exhibit distinct HRF nonlinearities compared to the neocortex (73). The primary goal of this study was to evaluate the feasibility and validity of an HC-tailored VASO protocol, both of which are supported by our findings. Nevertheless, future studies with larger cohorts and optimized modeling of HRF will be necessary to achieve a higher statistical power.

In conclusion, this study demonstrates that mesoscale VASO, when carefully optimized, is a viable tool for resolving laminar activation profiles in human HC. The protocol’s reproducibility across 7T platforms and sensitivity to task-evoked signals establish a strong foundation for future research to explore the depth-dependent properties of the HC subfields during complex cognitive functions and their alterations in neurodegenerative diseases.

## Data and code availability

Anonymized data have been deposited on Zenodo and will be publicly available upon manuscript acceptance. All original codes are provided in this GitHub repository: https://github.com/khazarAhmadi/VASO_HC.

## Authors contribution

K.A: Conceptualization, Methodology, Data acquisition, Formal analysis, Visualization, Writing – original draft, Writing – review and editing; S.S: Methodology, Writing – review and editing; S.K: Formal analysis, Writing – review and editing; A.B: Methodology, Writing – review and editing; P.B: Conceptualization, Writing – review and editing; N.A: Conceptualization, Writing – review and editing, Funding acquisition; L.H: Conceptualization, Methodology, Data acquisition, Supervision, Writing – review and editing, Funding acquisition.

## Acknowledgements

This project was supported by the European Research Council (grant no. 864164), Mercator Research Center Ruhr (grant no. Ko-2021-0010) and the NIMH/NINDS Intramural Research Program (#ZIAMH002783, #ZICMH002884). We thank Rüdiger Stirnberg from DZNE Bonn for the VASO 3D-EPI sequence code used here. We also thank Daniel Handwerker for institutional advice on ramping up the project. Special thanks to A. Tyler Morgan for serving as the scanner operator for one of the scan sessions and Marly Rubin, for her assistance with participant recruitment for some sessions. We would also like to acknowledge Viktor Pfaffenrot for valuable discussions on the analysis and interpretation of the results. Lastly, we thank Jordan DeKraker for encouraging us to undertake this study and to use HippUnfold tools.

For parallel initiatives involving layer-fMRI VASO in the hippocampus, please refer to a recent study from the Berkeley group at the Next-Gen 7T scanner (see ref.68).

## Supplementary Materials

**Supplementary Table 1.**
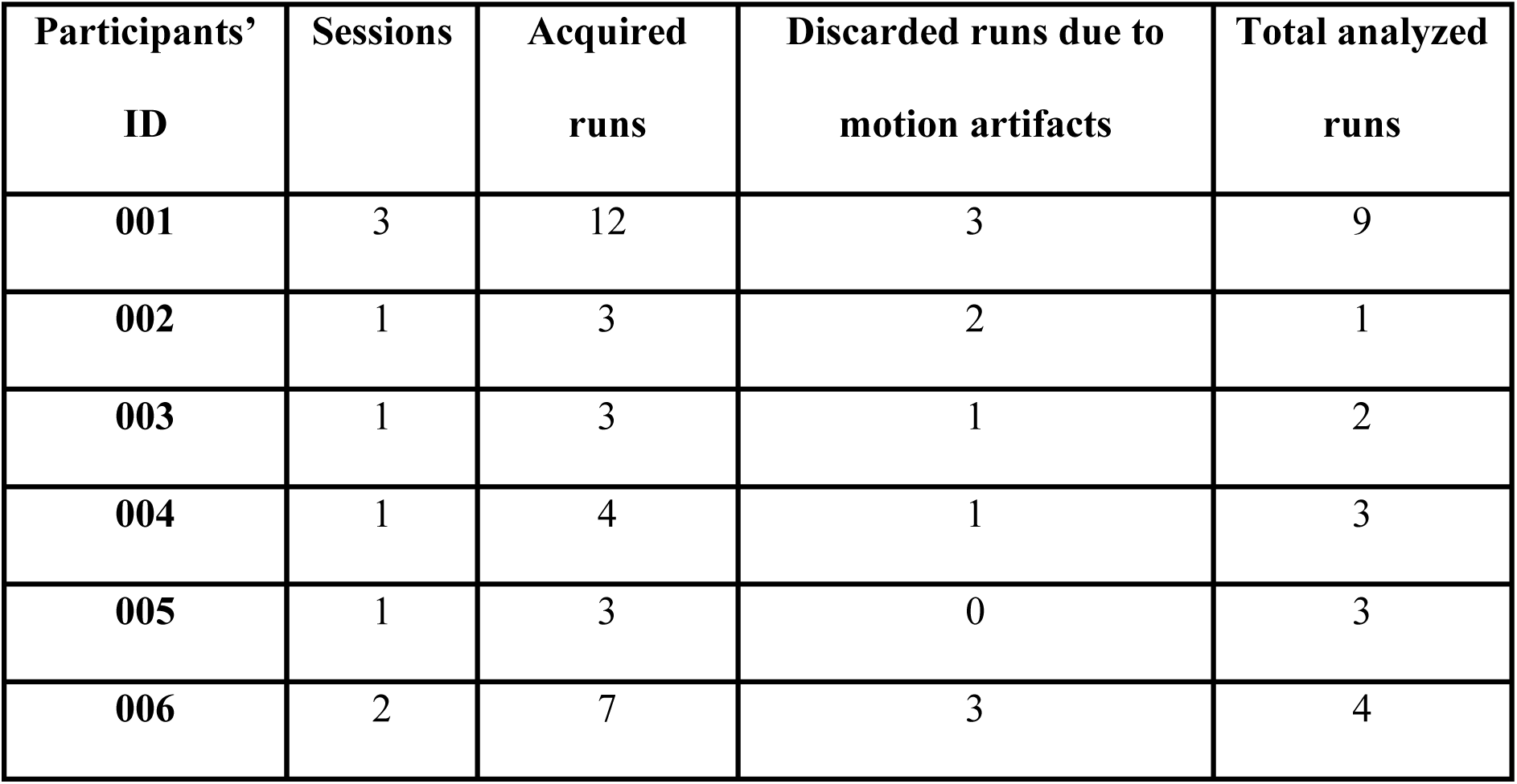
Overview of the number of acquired sessions and functional runs from six participants whose data were analyzed for part 2 of the current study, focused on validating the HC-tailored VASO sequence using an autobiographical memory task.

| **Participants’**<br>**ID** | **Sessions** | **Acquired**<br>**runs** | **Discarded runs due to**<br>**motion artifacts** | **Total analyzed**<br>**runs** |
| --- | --- | --- | --- | --- |
| **001** | 3 | 12 | 3 | 9 |
| **002** | 1 | 3 | 2 | 1 |
| **003** | 1 | 3 | 1 | 2 |
| **004** | 1 | 4 | 1 | 3 |
| **005** | 1 | 3 | 0 | 3 |
| **006** | 2 | 7 | 3 | 4 |

**Supplementary Figure 1.**
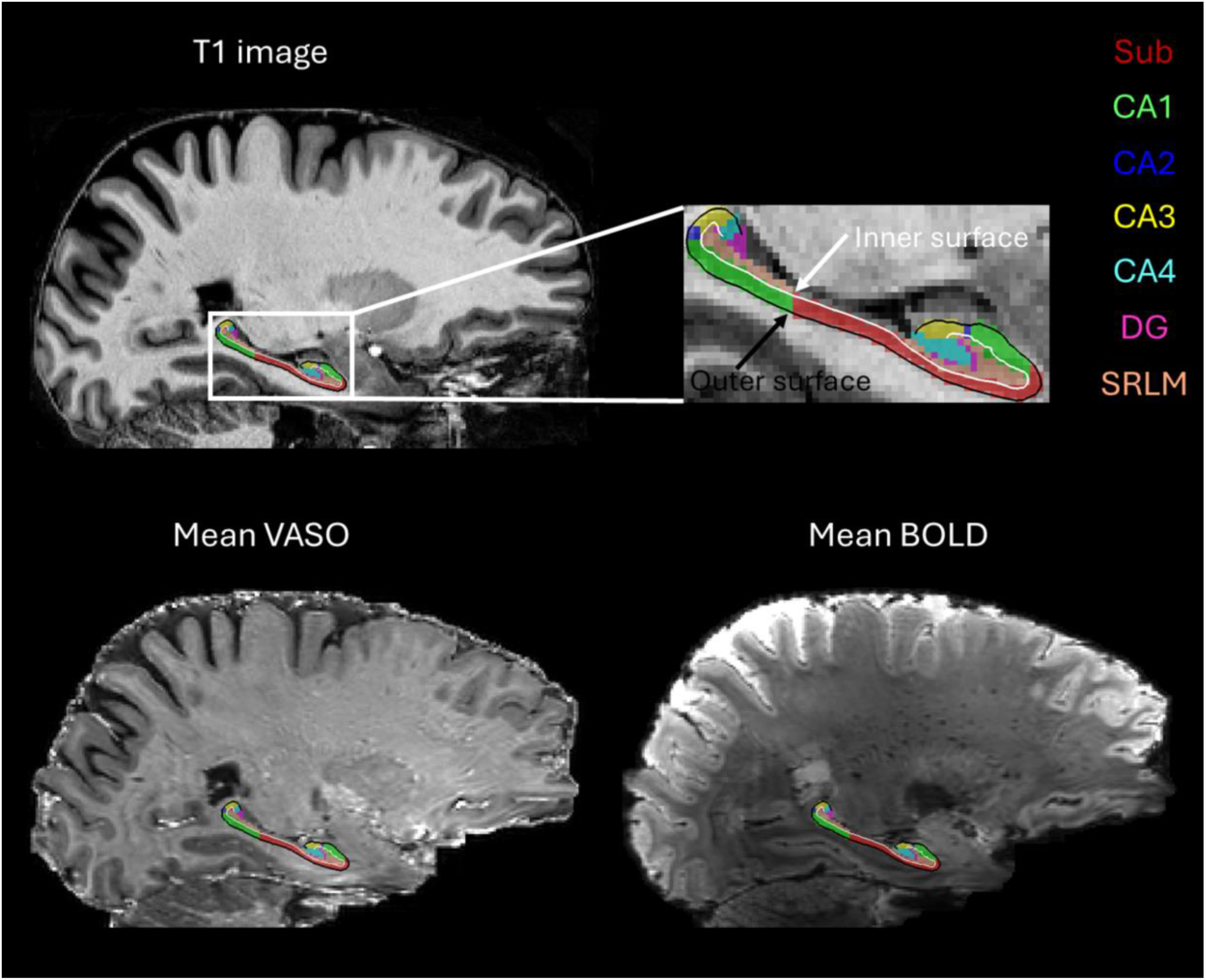
Hippocampal subfield segmentation and surface boundary delineation overlaid on a T1-weighted image from one participant. The bottom panel is the same as above but displayed on co-registered mean VASO and BOLD images. Subfield labels and inner/outer surfaces were generated using HippUnfold. Sub = subiculum, CA1-CA4 = cornu ammonis areas 1 to 4. DG = dentate gyrus, SRLM = stratum radiatum lacunosom moleculare.

**Supplementary Figure 2.**
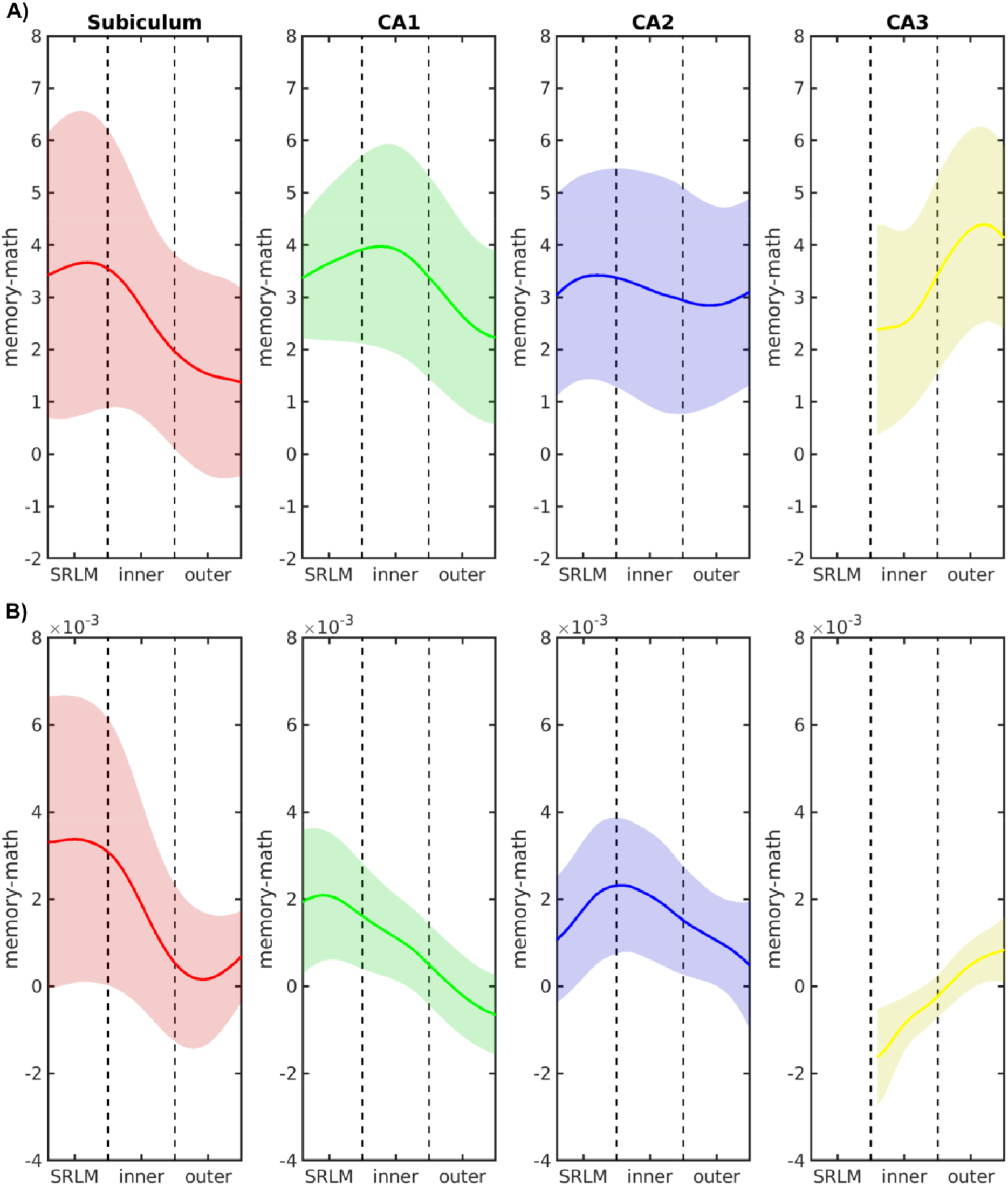
Laminar profiles of HC subfields for memory vs math contrast (non-transformed). **A)** BOLD signal modulations across the depths of subfields **B)** VASO signal modulations. The laminar profiles of both BOLD and VASO are largely consistent with the patterns observed in z-transformed memory vs. math contrast (see Figure.5).

